# Fine-scale temporal analysis of genotype-dependent mortality at settlement in the Pacific oyster *Crassostrea gigas*

**DOI:** 10.1101/084616

**Authors:** Louis V Plough

## Abstract

Settlement and metamorphosis mark a critical transition in the life cycle of marine invertebrates, during which larvae undergo substantial morphological, sensory, and genetic changes. High mortality during or after metamorphosis is commonly observed in both wild and hatchery settings, however, the underlying causes of this mortality remain poorly understood. Previous pair-crossing experiments with the Pacific oyster, *Crassostrea gigas* showed that substantial genotype-dependent mortality (GDM) occurs around metamorphosis, but, owing to sparse temporal sampling, it remains unknown whether mortality occurs just before, during, or after settlement. In this laboratory study, microsatellite marker segregation ratios were followed daily throughout the settlement and metamorphosis of an inbred, F_2_ cross of the Pacific oyster to examine the fine-scale patterns of GDM in larvae and spat. Genetic control of settlement timing was also examined using a quantitative trait locus (QTL) mapping approach. Settlement occurred over nine days (day 18 to day 27 post-fertilization) with 68% of individuals settling on an early (day 19) and a late (day 24) time point. Tracking the survival of spat for 40 days after initial settlement revealed almost no post-settlement mortality. Temporal analysis revealed that three of 11 loci exhibited segregation distortion at metamorphosis, one of which (*Cg205*) was followed throughout settlement. Alternative temporal patterns of selection against each homozygote at *Cg205* suggest possible defects in both the competency pathway (inability to initiate metamorphosis) and the morphogenesis pathway (mortality during the metamorphic transition). QTL mapping of settlement timing identified three individual and one epistatic QTL (29% of the variance explained), however, two of these loci were closely linked to markers exhibiting GDM at metamorphosis, thus making it difficult to distinguish between genetic variance in settlement timing and differential mortality early or late in settlement. Overall, results from this study highlight the complex temporal patterns of viability selection during metamorphosis and show that endogenous mortality during the larval-juvenile transition appears to be focused during or just prior to metamorphosis. Fine-scale experimental analysis of settlement can reveal important genetic insights into larval settlement behavior and the sources of larval mortality, and future studies should be able to further dissect the functional targets of selection during metamorphosis.

## 1 Introduction

The period of settlement and metamorphosis is critical in the life cycle of marine invertebrates. It is the final developmental hurdle to successful recruitment, wherein larvae go through the dramatic ecological and biological transformation from a free-swimming planktonic larval form to a sessile benthic juvenile. The successful completion of this process has profound implications for the evolution and ecology of species in the marine environment (Gaines and Roughgarden 1985, Hunt and Shiebling 1997, Roughgarden *et al.* 1988, Rodriquez *et al.* 1993). In marine mollusks, metamorphosis is well described, characterized by the complete rearrangement of the body plan, coinciding with the loss of larval features, such as the velum, and the emergence of juvenile or adult characteristics, such as the gills (*e.g.* Cole 1938, Hickman and Gruffyd 1971, Bonar 1976). The process of settlement and metamorphosis comprises two distinct phases: 1) the attainment of ‘competency’, the developmental capacity to respond to appropriate settlement cues and to exhibit settlement behavior, and 2) the morphogenetic transformation that occurs when larvae attach to the substratum and complete metamorphosis (e.g. Degnan and Morse 1995). The role and variety of chemical cues in the induction of metamorphosis, the developmental attainment of competency, and the consequences of delayed metamorphosis have been the subjects of much research in the field of larval biology and ecology (reviewed by Crisp 1974, Degnan and Morse 1995, Pechenik 1990, Rodriguez *et al.* 1993, Hadfield 1986, Pawlik 1992, Jackson *et al.* 2002).

Developmental and gene-expression studies in marine molluscs have revealed the complex nature of the metamorphic transition, showing that the competency and morphogenetic pathways display unique gene-expression profiles and are regulated independently (*e.g.* Degnan and Morse 1995, Jackson *et al.* 2005, Jackson *et al*. 2007, Williams *et al.* 2009). Patterns of increased gene expression late in the larval stages co-occur with the accumulation and deployment of a threshold level of receptors and signal transducers, which can then respond to inducible chemical cues that initiate metamorphosis (Degnan and Morse 1995, Jackson *et al.* 2005, Heylund and Moroz 2006). A suite of genes control an anticipatory, ‘competency’ pathway, which begins to form juvenile structures prior to metamorphosis (*e.g.* digestive and shell formation pathways; Jackson *et al*. 2007), and then the morphogenetic transformation is dictated by the up-regulation of genes related to apoptosis, cell cycling, protein synthesis, and calcium flux pathways, among others (the ‘morphogenetic’ pathway; Jackson *et al.* 2005, Jackson and Degnan 2006, Jackson *et al.* 2007, Williams *et al.* 2009). Though there appear to be clear patterns of gene classes expressed across invertebrate taxa during competency and metamorphosis (*e.g.* Heylund and Moroz 2006), most studies have focused on only a few marine gastropods (*Aplysia* and *Haliotus* spp.) and ascidians (*e.g.* Degnan *et al.* 1997, Eri *et al*. 1999, Kawashima 2005, Jacobs *et al.* 2006), all of which have lecithotrophic larvae. Less is known about genetic processes at metamorphosis in broadcast spawning marine invertebrates, such as marine bivalves, with planktotrophic larvae.

Metamorphosis in marine invertebrates, and marine bivalves in particular, is accompanied by substantial mortality. High rates of mortality in newly settled juveniles have been observed for a wide variety of invertebrate species, and, generally, survival curves of new settlers are type III: mortality is initially high but decreases rapidly after the first few days or weeks and then levels off (*e.g.* Rodriguez *et al*. 1993, reviewed by Gosselin and Qian 1997, Hunt and Sheibling 1997). Early mortality in invertebrates has primarily been estimated in field studies that examine rates and patterns of mortality after settlement (*e.g.*, Rodriguez *et al.* 1993, Gosselin and Qian 1996, Hunt and Shiebling 1997); however, many of these studies are only able to measure “recruitment”, *i.e.*, the survival of a population to a certain time point after settlement, because monitoring mortality during settlement is difficult, if not impossible, for many species (*e.g.* Keough and Downes 1982). Few studies accurately measure larval supply, metamorphosis, and early and late post-settlement periods within the same experiment, thus, a comprehensive understanding of when mortality occurs during settlement and metamorphosis is lacking.

Experimental laboratory studies allow for more detailed analysis of mortality during settlement in marine invertebrates, but few such studies exist. Jones and Jones (1983) found that only a small fraction (10-30%) of pre-metamorphic oyster larvae successfully complete metamorphosis in laboratory conditions. Haws *et al.* (1993) also noted that substantial mortality occurred during the metamorphic transition in their detailed examination of biochemical and physiological changes during metamorphosis of the Pacific oyster *Crassostrea gigas*. Culture experiments have shown significantly greater post-metamorphic survival associated with improved diet during the larval stages, suggesting that endogenous processes related to energy acquisition prior to metamorphosis are important (*e.g.* Gallager et al 1986, Gallager and Mann 1986, Helm *et al.* 1991, Coutteau *et al.* 1994, Pernet and Tremblay 2004). Recent experimental genetic work with the Pacific oyster has confirmed high mortality at metamorphosis, finding that half of the deleterious recessive mutations uncovered in inbred crosses are expressed at this stage of the life cycle (Plough and Hedgecock 2011). Plough and Hedgecock (2011) determined the stage-specific timing of genotype-dependent mortality by following the temporal onset of genotype deficiencies (selection against specific genotypes) in daily larval samples, a post settlement sample, and an adult sample. The finding of substantial, genetic mortality during settlement highlights the important physiological and developmental changes associated with this transition, however, there were no temporal genetic data collected during metamorphosis in Plough and Hedgecock (2011), and thus the fine scale temporal patterns of selection remain unknown.

In the current study, temporal genetic sampling was focused on the period of settlement and metamorphosis to determine the detailed timing of genotype dependent mortality during metamorphosis. An inbred, F_2_ family was allowed to set naturally on adult shell, and daily samples were taken of spat (recently settled juvenile oysters) and larvae still residing in the water column, throughout the settlement period. Utilizing a similar experimental design to Plough and Hedgecock (2011), mutations affecting metamorphosis were first identified by the specific developmental timing of segregation distortion in linked genetic markers: *i.e.* those markers that were observed to be distorted in juvenile samples (day 60, post-settlement) but were in Mendelian proportions in pre-metamorphic, late larval samples were identified as candidates. Then, the segregation ratios and genotype deficiencies of these metamorphosis-associated markers were followed through late larval and spat populations, to examine the temporal patterns of selection during and post-metamorphosis. Three hypotheses are proposed to explain how selection and mortality might proceed: 1) larvae are able to advance through metamorphosis and settle, but die soon after, which can be detected by high mortality of juveniles after settlement and changes in segregation ratios in early vs. late juvenile samples; 2) larvae begin metamorphosis, but die sometime during this transition, thus, affected genotypes are present in the water column at time *i* (***t***_*i*_), but at ***t***_i+1_, they are missing from both the spat and larval pool; and 3) individuals with affected genotypes are not able to begin metamorphosis and are thus are deficient in the spat population but remain and perhaps accumulate in the larval population, because settlement is delayed or fails to occur altogether. Finally, early vs. late settling spat were genotyped to determine if there were markers or regions of the genome associated with variation in settlement timing that were independent of those causing mortality at metamorphosis. The detailed sampling and genotyping strategy carried out in this experiment sheds light on the fine-scale timing of genotype-dependent mortality at settlement, and determines the extent to which competency, morphogenesis, or post settlement processes are affected by the expression of genetic load.

## 2. Materials and Methods

### 2.1. Crosses and culturing methods

Inbred lines *51* and *35* were derived from a naturalized population of *C. gigas* in Dabob Bay, WA, with initial families made from pair crosses of wild individuals in 1996 (Hedgecock and Davis 2007). These lines were inbred (full-sib mating) for four generations leading up to the F_1_ hybrid cross that was made in 2007. In 2009, the experimental F_2_ family (*f* = 0.397) was created by mating a pair of male and female full-siblings from the 2007 *51*×*35* F_1_ hybrid cross. Crosses were performed at the University of Southern California (USC), Wrigley Marine Science Center (WMSC) on Catalina Island, CA. Pedigrees of parents were verified with microsatellite DNA markers (e.g. Hedgecock and Davis 2007)

The cross was performed by stripping gametes from a single male and a single female and combining the eggs and sperm in a two-liter beaker of fresh seawater for fertilization (Breese and Malouf 1975, Hedgecock and Davis 2007). After a one-hour incubation, one million zygotes were stocked in a 200-l vessel (5 larvae·ml^-1^) with fresh sea water. Starting at 24 h post-fertilization, larvae were fed a diet of *Isochrysis galbana* (Tahitian Isolate) every two days, at a starting concentration of 30,000 cells·ml^-1^, which was increased, as larvae grew, following standard larval rearing protocols for the Pacific oyster (Breese and Malouf 1975, Hedgecock *et al.* 1996, Plough 2012).

### 2.2. Deployment of shell and sampling protocol

At day 18, a substantial number of larvae had developed eye spots and were displaying probing with the larval foot, both characteristic of settlement behavior (Bonar 1976, Kennedy *et al.* 1996). Cured adult shell was deployed on a cylindrical harness with a bottom of Vexar® mesh, as a substrate for natural settlement (Fig. 1a). Adult shell was placed with the nacreous side facing both up and down and with shells overlapping to form random 3-D structures, in order to give larvae a number of different substrates and angles to settle on regardless of their orientation to light or gravity, which can affect settlement behavior (Kennedy *et al.* 1996, Baker 1997, Baker and Mann 1998). Starting in the afternoon on day 18, and at about the same time each day thereafter, the harness was carefully pulled up, and the surface of shells were sprayed vigorously with seawater to return any probing or non-settled larvae back to the 200-l vessel and the larval pool. Larvae were filtered out of the 200-l vessel and concentrated for survival estimates each day during settlement. Survival was calculated by counting the number of larvae in four to six random sub-samples of known volume (50-100 µl) from homogeneously mixed, concentrated cultures (500-1000 ml). These counts were then multiplied by the inverse of the fraction of the subsample volume to the total volume 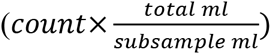, and averaged across replicates to estimate the mean number of larvae remaining (surviving) each day. For each daily estimate of settlement, spat were counted shell by shell, under a dissecting microscope at 8-48×, until no more could be found. This involved inspecting both sides of the shell under a range of magnifications and orientations. New shells that had been incubating for 24 hours in fresh, filtered seawater, were then placed on the harness, and lowered back into the 200-l tank, which had been rinsed and refilled with fresh seawater and algae. Finally, the counted larvae were placed back into the tank. This process was repeated each day until no larvae remained in the water column. From the counts of larvae, *L*, at time *i* (***t***_*i*_) and ***t***_*i+1*_, and the count of settlers, *S*, at ***t***_*i+1,*_ larval mortality (M) between two adjacent days was calculated as: *M* = *L_i_* (*L*_*i*+1_ + *S*_*i*+1_). Shells that had spat settling on a particular day were then collected and placed in a Vexar® mesh bag in the upwelling nursery system and grown out until day 60 to increase the amount of tissue for DNA extraction (Fig 1b). Spat were fed live *Isochrysis galbana* and Shellfish Diet 1800 (Reed Mariculture, Campbell, CA) every 6 six hours.

**Figure 1.**
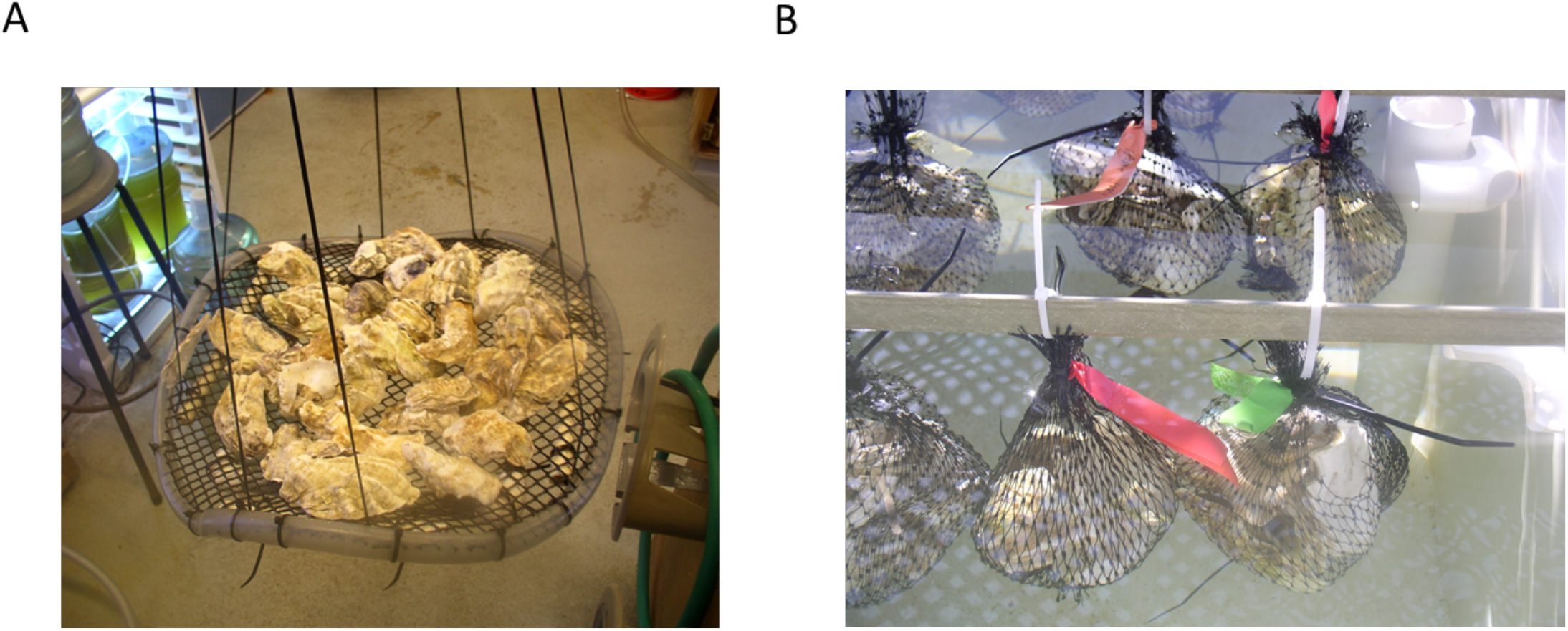
Pictures of shell deployment and bags with spat for grow-out. The collection of shells deployed each day on the Vexar® harness (A), and shells with spat separated in bags in the nursery system for grow-out (B).

To determine if mortality occurred post-settlement (after the day on which individuals were recorded as settling), a subset of metamorphosed spat from day 19, 21, and 24, were followed to day 60 to determine survival through the early juvenile stage. This was done by marking an area around 50 (live) spat on two shells from each of the three days, and then counting the number of live spat on these same, marked shell areas at day 60.

### 2.3. DNA extraction, PCR, and electrophoresis

At 60 days, individual spat were scraped off shells and placed directly in Qiagen tissue lysis buffer (Qiagen, Valencia CA) in individual tubes and stored at -80ºC until extraction using the Qiagen DNeasy blood and tissue kit. Live larvae were placed in small volumes of sterilized water with a drop of 70% ethanol to retard their movement; then, they were pipetted individually into 96-well PCR plates containing extraction buffer consisting of 1× PCR buffer (Promega, Madison,Wisconsin), 0.2 mM EDTA, 1.0 µg/µl proteinase K (Shelton Scientific, Connecticut) and purified H_2_O. Plates were frozen at -80ºC until extraction (Plough and Hedgecock 2011). Larvae were extracted in 40 µl volumes in a 96-well thermalcycler (BioRad Tetrad, Hercules, CA), held at 56ºC for three hours, followed by 15 minutes at 95ºC. Parent tissue was stored in 70% ethanol at 4ºC until extraction and DNA was extracted from 10-25 mg of tissue, using either the Qiagen DNeasy animal tissue kit or the Gentra Puregene tissue kit (Qiagen, Valencia CA), following the manufacturer’s protocols.

Eighty-four microsatellite markers cloned from the Pacific oyster were tested in this study (Magoulas *et al.* 1998, McGoldrick *et al.* 2000, Huvet 2000, Li *et al.* 2003, Sekino *et al.* 2003, Yamtich *et al.* 2005, Yu and Li 2007, Wang *et al.* 2007). Genotyping of microsatellites was carried out as described previously (Plough and Hedgecock 2011, Plough 2012) and 46 markers were found to be informative, all of which are on published linkage maps (Hubert and Hedgecock 2004, Hubert *et al.* 2009, Plough and Hedgecock 2011). From these 46 markers, a subset of 24 were chosen for the QTL mapping analysis (see below), with the aim of having at least two markers on each of the 10 linkage groups, thereby maximizing genomic coverage with the fewest markers.

### 2.4. Genetic analysis of mortality during settlement

To identify markers that became distorted during settlement/metamorphosis, markers were, first, tested for significant deviations from expected Mendelian ratios in a sample of 60-day-old spat (*n* = 192) made up of a pooled sample of individuals that settled either on day 19 or day 24. These two days were chosen because they represented the bulk of the settlement (68% of spat settled on day 19 or day 24; Fig. 2), making the pooled sample fairly representative of the total settled population that would have been sampled at day 60 without regard to settlement timing. Markers were only considered for analysis during metamorphosis if they exhibited segregation distortion in the end-point, day 60 pooled spat sample. Once markers exhibiting segregation distortion in the pooled spat sample were identified, they were genotyped in day-18 larval samples to identify markers that exhibited segregation distortion (selection) before metamorphosis and thus were not candidates of genotype dependent mortality at metamorphosis. Markers that exhibited Mendelian segregation ratios just before metamorphosis (day 18) but showed distorted segregation ratios in the day-60 spat sample were then genotyped in larvae and spat from each day throughout settlement and metamorphosis.

**Figure 2.**
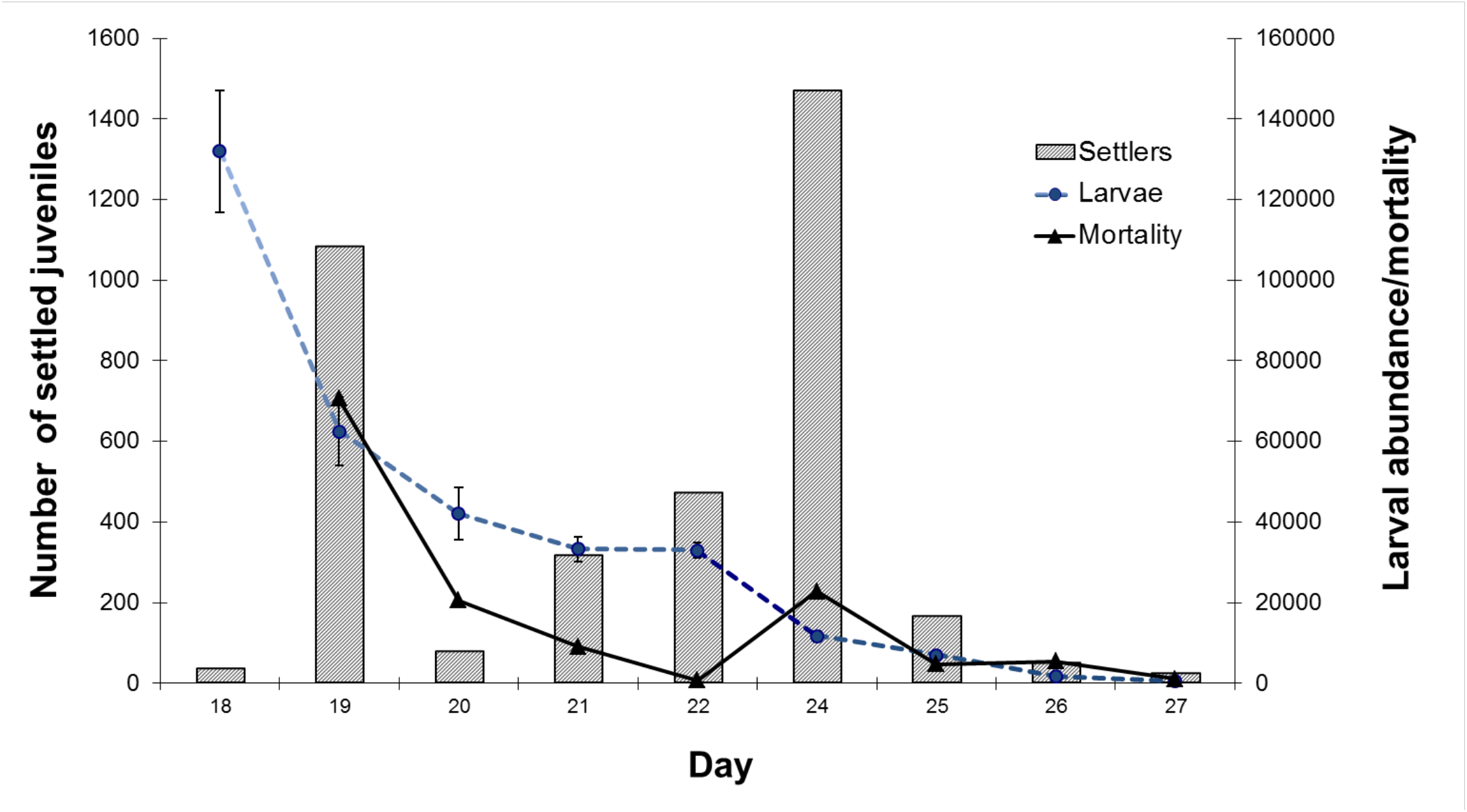
Settlement and Mortality data. Mortality is calculated as the difference in the abundance of larvae between days, accounting for the number new settlers. Error bars on the larval abundance estimates represent one standard error of the mean from volumetric counts.

### 2.5. Genetic analysis of settlement timing

Spat settling on day 19 and day 24 (see Fig. 2) were used to examine possible genetic differences between early and late settling individuals. For an initial, single marker analysis, 96 spat from the early and the late settlement samples were genotyped at 24 loci and tested for heterogeneity between the two settlement samples with contingency chi-square tests using the program ‘Chirxc’ (Zaykin and Pudovkin 1992). For the genome-wide analysis of variation in settlement timing, linkage maps were first constructed with day 19 and day 24 spat samples pooled (*n* = 192), using the CP (cross pollinator) population type in JoinMap 3.0 (VanOoijen and Voorrips 2001). The Kosambi mapping function with a minimum likelihood of the odds (LOD) score of 2.0 was used for linkage group assignments, though most linkage groups had LOD scores well above 4.0. Phase was determined by JoinMap 3.0, from the frequencies of parental and recombinant types. Deviations from Mendelian segregation ratios may affect linkage mapping (Zhu *et al.* 2007), so marker order and distances were compared with previously published linkage maps, which were constructed using larval samples with little segregation distortion (e.g. Hubert and Hedgecock 2004). If markers that should have mapped, failed to map, locations were taken from published maps (Hubert and Hedgecock 2004, Hubert *et al.* 2009, Plough and Hedgecock 2011). Genomic coverage was estimated with the equation *GC* = 1 084616 *e*^−2\**dn/L*^, where *d* = mean inter-marker distance and *n* = total number of markers assigned to linkage groups, and *L* is the estimated map length (Bishop *et al.* 1983).

QTL analysis was performed in R/qtl (v.1.1; Broman *et al.* 2003), under a binary trait model, (early vs. late settlement, day 19 vs. day 24), using the outcross settings. A one-dimensional scan for single QTL was first performed (‘scan-one’ module) and 5% significance thresholds at the genome-wide and chromosome-wide level were calculated with 1000 permutations. Next, genetic interaction was tested at every position in the genome, using a two-QTL model (‘scan-two’ module; significance adjusted with 1000 permutations). Once significant QTL were identified, they were fit into an overall multiple regression ANOVA model, using a backwards selection procedure to find the best model with lowest model comparison criterion.

## 3. Results

### 3.1. Observed settlement and mortality

Larvae showed the characteristic signs of settlement behavior (*e.g.* eyespots and probing foot) on day 17 post-fertilization and adult shells were deployed starting on day 18. Around 30 larvae had already settled on day 17, sticking to the bottom of the poly-carbonate tank in the absence of adult shell; these were not sampled. On the initial day of sampling, day 18, the number of surviving larvae was 132,000 ± 7,500 (from a starting stock of 1,000,000 embryos), which represents 13.2% survival to the end of the larval stage (Fig. 2). Settlement occurred over nine days, from day 18 to day 27, with settlement peaks at day 19 (1,084 counted spat) and day 24 (1,484 spat), indicating that settlement in this family followed a bi-modal distribution. In total, 3,776 settlers were counted, representing 0.37 % survival to the spat stage over the 60 days of the experiment. Larval abundance appeared to drop sharply on the days when a substantial number of settlers were recorded (*e.g.* day 19 and day 24) and a corresponding spike in larval mortality was observed, despite the correction for settled spat (Fig. 2). After day 24, the abundance of larvae steadily declined, despite few additional spat setting. Mortality of new settlers followed for approximately 40 days after initial settlement (examined at day 60) was minimal, with estimates of 2%, 0% and 2% for settlers on day 19, 21, and 24, respectively. In general, very few dead or empty shells (spat) from any settlement day were observed. Because very little post-settlement mortality was observed, segregation ratios at markers were not followed from initial settlement samples to day 60 settlement samples.

### 3.2. Segregation analyses and segregation distortion during settlement

Eighty-four microsatellite markers were tested, yielding 46 informative markers in the parents of the experimental progeny. A subset of 24 markers, comprising at least two loci on each of the 10 Pacific oyster linkage groups (LG), were tested for distortion of segregation ratios in spat settling on day 19 (*n* = 96) and spat settling on day 24 (*n* = 96), which were harvested (sampled) at 60 days post fertilization. None of the three informative markers on LG 2 could be scored, so these markers and this linkage group were not included in the analysis, leaving 21 markers for further analysis. Significantly distorted segregation ratios at the α = 0.05 level were found at 13 of 21 (62%) markers (Table 1). These distortions were mainly caused by deficiencies of homozygous genotypes or heterozygote carriers of parentally shared, presumably identical-by-descent, deleterious alleles. Markers with distorted segregation ratios were distributed broadly across the genome, on eight out of the nine linkage groups analyzed. Genotyping all but two (*Cg162* and *Cg156*) post-settlement distorted markers in the day-18 larval sample revealed that markers *Cg109*, *Cg205*, and *Cg175* on LG 4, LG 6, and LG 8, respectively, exhibited Mendelian segregation ratios (three of 11 tested; Table 1), indicating that deleterious alleles linked to these markers were first expressed during or just after metamorphosis.

**Table 1.**
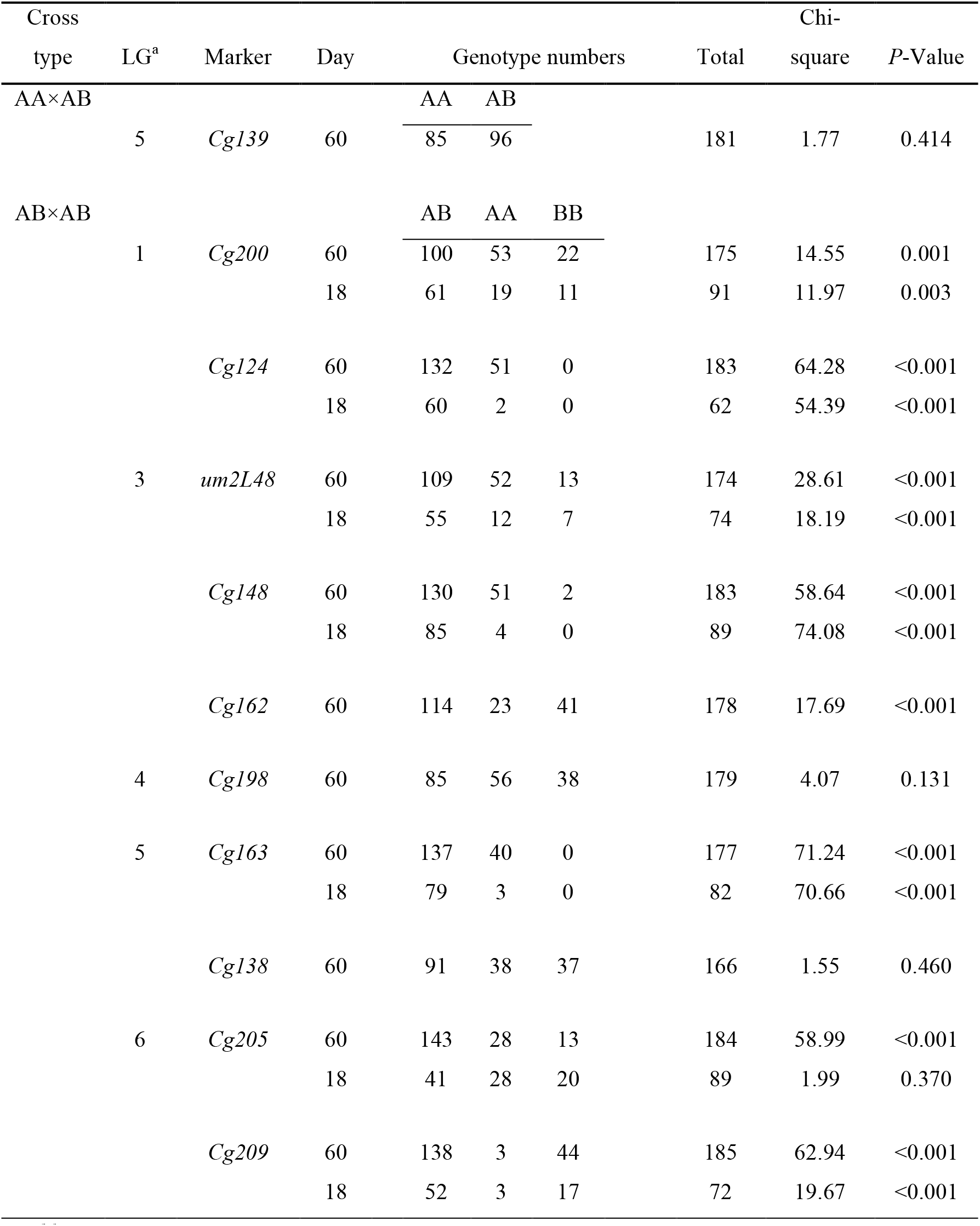
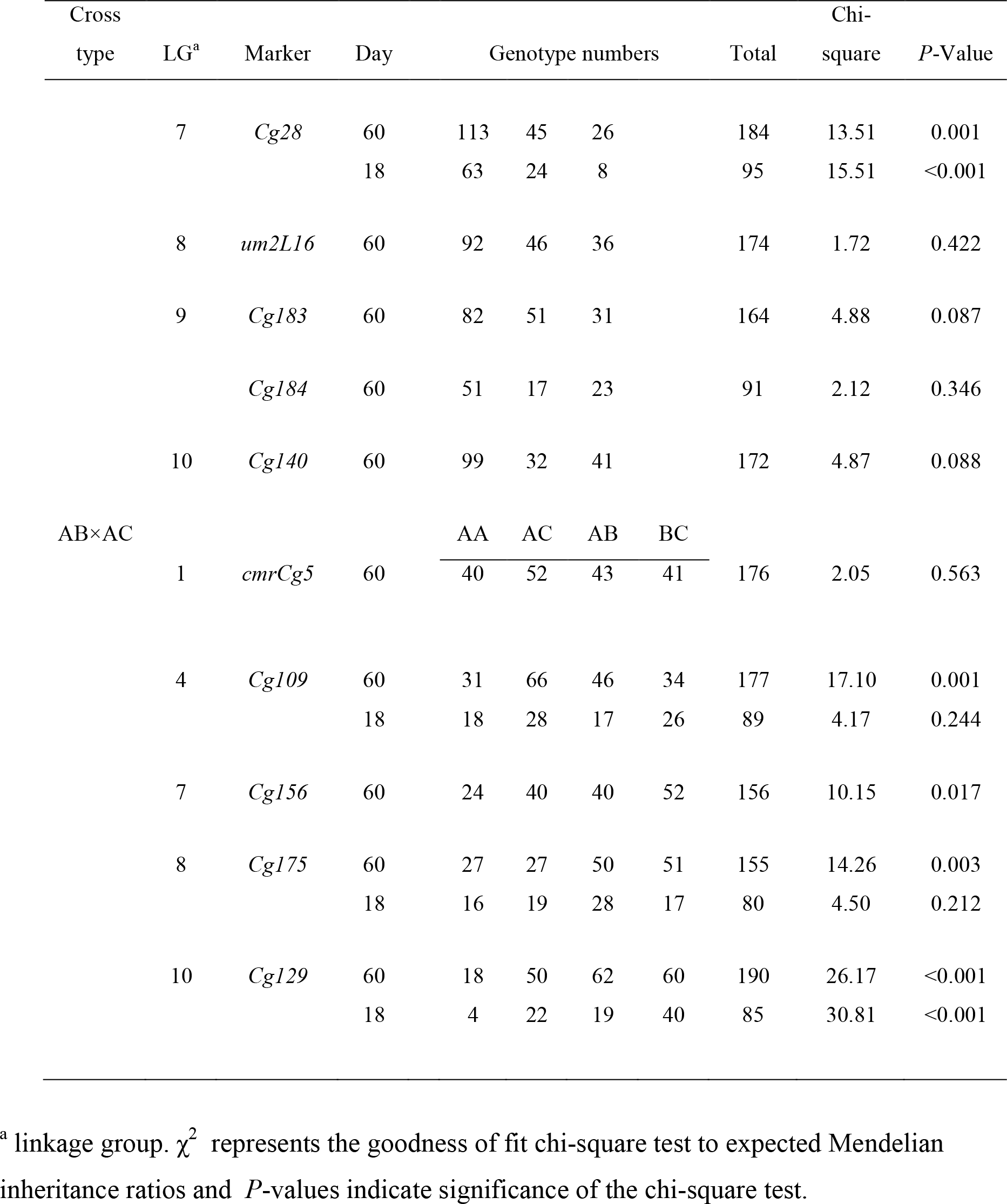
Marker segregation data for day 18 larvae and day 60 spat

| Cross<br>type | LG<sup>a</sup> | Marker | Day | Genotype numbers |  |  | Total | Chi-square | P-Value |
| --- | --- | --- | --- | --- | --- | --- | --- | --- | --- |
| AA×AB |  |  |  | AA | AB |  |  |  |  |
|  | 5 | *Cg139* | 60 | 85 | 96 |  | 181 | 1.77 | 0.414 |
| AB×AB |  |  |  | AB | AA | BB |  |  |  |
|  | 1 | *Cg200* | 60 | 100 | 53 | 22 | 175 | 14.55 | 0.001 |
|  |  |  | 18 | 61 | 19 | 11 | 91 | 11.97 | 0.003 |
|  |  | *Cg124* | 60 | 132 | 51 | 0 | 183 | 64.28 | <0.001 |
|  |  |  | 18 | 60 | 2 | 0 | 62 | 54.39 | <0.001 |
|  | 3 | *um2L48* | 60 | 109 | 52 | 13 | 174 | 28.61 | <0.001 |
|  |  |  | 18 | 55 | 12 | 7 | 74 | 18.19 | <0.001 |
|  |  | *Cg148* | 60 | 130 | 51 | 2 | 183 | 58.64 | <0.001 |
|  |  |  | 18 | 85 | 4 | 0 | 89 | 74.08 | <0.001 |
|  |  | *Cg162* | 60 | 114 | 23 | 41 | 178 | 17.69 | <0.001 |
|  | 4 | *Cg198* | 60 | 85 | 56 | 38 | 179 | 4.07 | 0.131 |
|  | 5 | *Cg163* | 60 | 137 | 40 | 0 | 177 | 71.24 | <0.001 |
|  |  |  | 18 | 79 | 3 | 0 | 82 | 70.66 | <0.001 |
|  |  | *Cg138* | 60 | 91 | 38 | 37 | 166 | 1.55 | 0.460 |
|  | 6 | *Cg205* | 60 | 143 | 28 | 13 | 184 | 58.99 | <0.001 |
|  |  |  | 18 | 41 | 28 | 20 | 89 | 1.99 | 0.370 |
|  |  | *Cg209* | 60 | 138 | 3 | 44 | 185 | 62.94 | <0.001 |
|  |  |  | 18 | 52 | 3 | 17 | 72 | 19.67 | <0.001 |

| Cross type | LG<sup>a</sup> | Marker | Day | Genotype numbers |  |  |  | Total | Chi-square | P-Value |
| --- | --- | --- | --- | --- | --- | --- | --- | --- | --- | --- |
|  | 7 | *Cg28* | 60 | 113 | 45 | 26 |  | 184 | 13.51 | 0.001 |
|  |  |  | 18 | 63 | 24 | 8 |  | 95 | 15.51 | <0.001 |
|  | 8 | *um2L16* | 60 | 92 | 46 | 36 |  | 174 | 1.72 | 0.422 |
|  | 9 | *Cg183* | 60 | 82 | 51 | 31 |  | 164 | 4.88 | 0.087 |
|  |  | *Cg184* | 60 | 51 | 17 | 23 |  | 91 | 2.12 | 0.346 |
|  | 10 | *Cg140* | 60 | 99 | 32 | 41 |  | 172 | 4.87 | 0.088 |
| AB×AC |  |  |  | AA | AC | AB | BC |  |  |  |
|  | 1 | *cmrCg5* | 60 | 40 | 52 | 43 | 41 | 176 | 2.05 | 0.563 |
|  | 4 | *Cg109* | 60 | 31 | 66 | 46 | 34 | 177 | 17.10 | 0.001 |
|  |  |  | 18 | 18 | 28 | 17 | 26 | 89 | 4.17 | 0.244 |
|  | 7 | *Cg156* | 60 | 24 | 40 | 40 | 52 | 156 | 10.15 | 0.017 |
|  | 8 | *Cg175* | 60 | 27 | 27 | 50 | 51 | 155 | 14.26 | 0.003 |
|  |  |  | 18 | 16 | 19 | 28 | 17 | 80 | 4.50 | 0.212 |
|  | 10 | *Cg129* | 60 | 18 | 50 | 62 | 60 | 190 | 26.17 | <0.001 |
|  |  |  | 18 | 4 | 22 | 19 | 40 | 85 | 30.81 | <0.001 |

<sup>a</sup> linkage group. $\chi^2$ represents the goodness of fit chi-square test to expected Mendelian inheritance ratios and *P*-values indicate significance of the chi-square test.

Difficulties in genotyping larval samples only allowed temporal analysis of *Cg205* after day 19. In the pooled spat sample for *Cg205* (post-metamorphosis, “cumulative” sample; Fig. 3), significant deficiencies in both homozygous genotypes were evident, suggesting that *Cg205* might be linked to two deleterious alleles in repulsion phase (see Plough and Hedgecock 2011). Starting with the inspection of genotypic data for day-19 spat, observed segregation ratios were highly distorted, with substantial deficiencies of the two homozygous genotypes (Table 2, Fig. 3). Segregation ratios in day-19 larvae did not deviate from their Mendelian expectation however, indicating that genotypes deficient in the spat sample were still present among larvae in the water column. At the next larval sample, day 21, segregation ratios were significantly distorted, but only the *AA* genotype was deficient; the *BB* genotype did not deviate from its expected 1:2 ratio, relative to the frequency of *AB*. At day 24, segregation ratios in the larval sample were dramatically different from the previous time point (day 21), with a much larger proportion of *BB* genotypes than expected (60% compared with the expected 25%), and significantly fewer *AB* genotypes, while the number (and relative frequency) of *AA* genotypes remained highly deficient (only five spat with *AA* genotypes were observed in both day 21 and day 24 larval samples; Fig. 3, Table 2). In the day-24 spat sample, both homozygous genotypes were again observed to be deficient, and segregation ratios were dramatically different from the day 19 sample, owing to a greater frequency of *BB* individuals settling (*P*<0.001, *r*×*c* contingency chi-square test; Zaykin and Pudovkin 1992). At the final larval time point, day 26, the *BB* genotype was in even greater excess (79% compared to the expected 25%), while the *AA* genotype was completely absent; the *AB* genotype was deficient at this time point as well (Table 2, Fig. 3). In the day-26 spat sample, a slight increase in *BB* genotypes was observed, but both the *BB* and *AA* genotypes were significantly deficient relative to Mendelian expectation, a result consistent with observed segregation data for the cumulative, total spat sample (days 19, 24, and 26 combined; Fig. 3). In summary, individuals with the *AA* genotype suffered relatively swift mortality early in settlement (day 19 to day 21) with few individuals surviving in the water column (larvae) or setting successfully after this point. In contrast, while *BB* individuals were also deficient in the spat samples relatively early in settlement (day 19), they were present in the larval pool throughout settlement, becoming the dominant genotype remaining in the water column late in the larval period, but never setting in appreciable numbers even in the last days of settlement. The differing temporal patterns of mortality and segregation distortion in the two homozygous genotypes suggest that *Cg205* is linked to two deleterious recessive mutations in repulsion phase, with very different patterns of selection (Plough and Hedgecock 2011).

**Table 2.** Genotype numbers for *Cg205* during settlement

| *Settlers Cg205* |  |  |  |  |  | *Larvae Cg205* |  |  |  |  |  |
| --- | --- | --- | --- | --- | --- | --- | --- | --- | --- | --- | --- |
| Day | *AB* | *AA* | *BB* | Chi-Square | *P*-Value | Day | *AB* | *AA* | *BB* | Chi-Square | *P*-Value |
| d18 | -- | -- | -- | -- | -- | d18 | 41 | 20 | 28 | 1.99 | 0.370 |
| d19 | 82 | 9 | 2 | 55.26 | <0.0001 | d19 | 36 | 14 | 17 | 0.64 | 0.725 |
| d21 | -- | -- | -- | -- | -- | d21 | 42 | 5 | 24 | 12.55 | 0.002 |
| d24 | 61 | 11 | 19 | 11.97 | 0.003 | d24 | 27 | 5 | 47 | 52.57 | <0.0001 |
| d26 | 30 | 3 | 7 | 10.80 | 0.005 | d26 | 7 | 0 | 27 | 54.65 | <0.0001 |
| **Cum.** | **173** | **23** | **28** | **86.50** | **<0.0001** |  |  |  |  |  |  |

Chi-square represents the goodness-of-fit chi-square test to expected Mendelian inheritance ratios.

**Figure 3.**
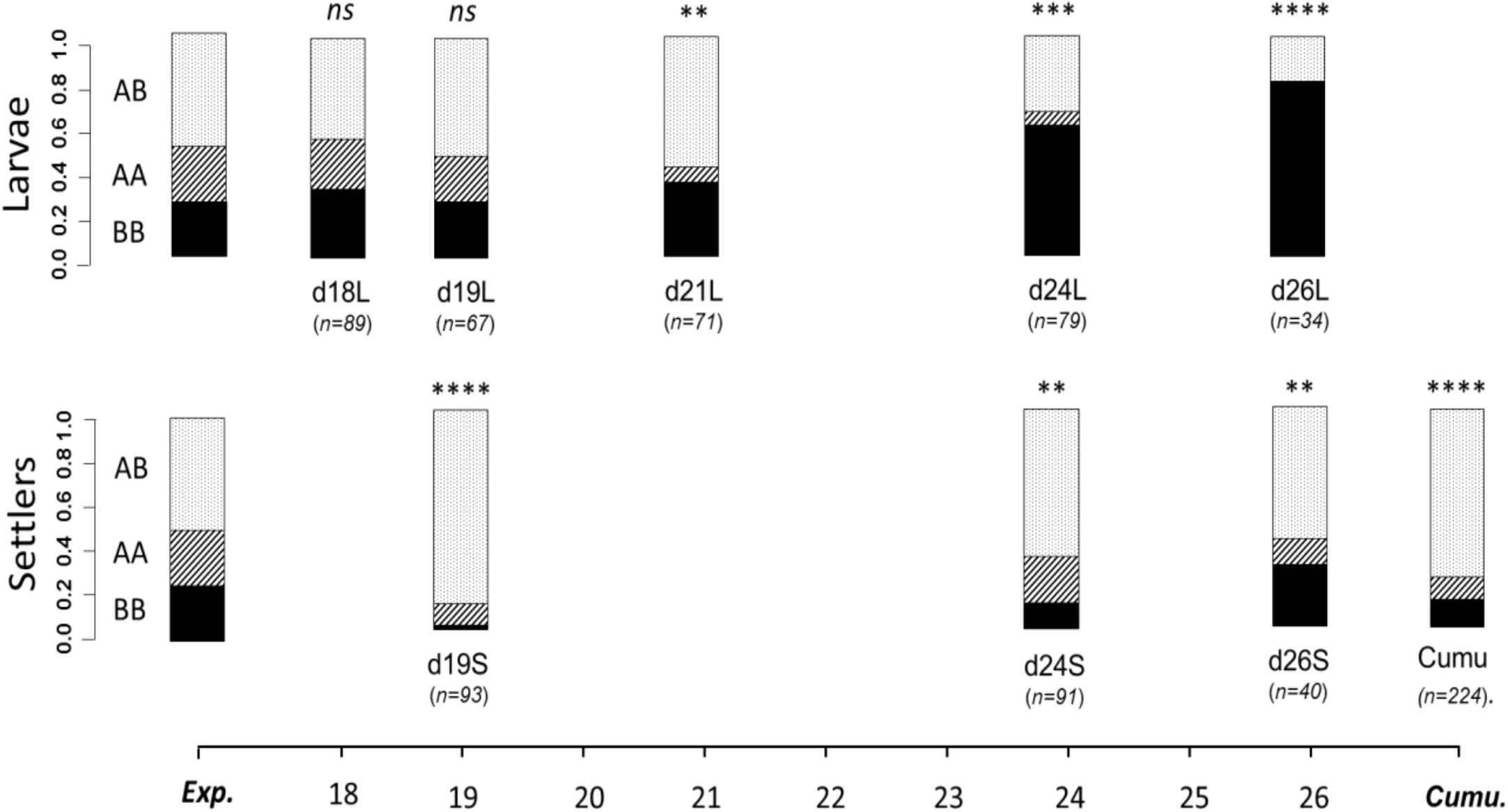
*Cg205* genotype proportions in larvae and spat during settlement. Asterisks denote the level of significance for the goodness-of-fit chi-square test (*cf.* Mendelian expectations of 2:1:1, *AB*:*AA*:*BB*) of genotype proportions at *Cg205* (* *P*<0.05, ** *P*<.01, *** *P*<0.001, **** *P*<0.0001); *ns* is non-significant. The cumulative (cumu) sample is the pooling of all spat collected during settlement (e.g. d19, d24, and d26).

### 3.3. QTL mapping and genetic analysis of settlement timing

Twenty-one markers were assigned to 8 of 10 linkage groups, with assignments as expected from previous crosses and studies (Hubert and Hedgecock 2004, Hubert *et al.* 2009, Plough and Hedgecock 2011), marker phase was successfully estimated within linkage groups. Linkage group two lacked informative markers, and map distances for the two markers on linkage group seven were obtained from previously published maps because markers were separated by more than 40 cM; phase was still successfully estimated from segregation data in this linkage group. Total map length was 369 cM, calculated by summing all inter-marker distances, with a mean inter-marker distance of 17.65 cM (maximum 42 cM; LG 7). Estimated genomic coverage for this map was 86% (Bishop *et al.* 1983). Single-marker analysis revealed that four of the 21 markers, *Cg205*, *Cg162*,*Cg140* and *Cg109*, had significant heterogeneity in genotype proportions between days 19 and 24, indicating that these markers might be associated with settlement timing (Table 3).

**Table 3.**
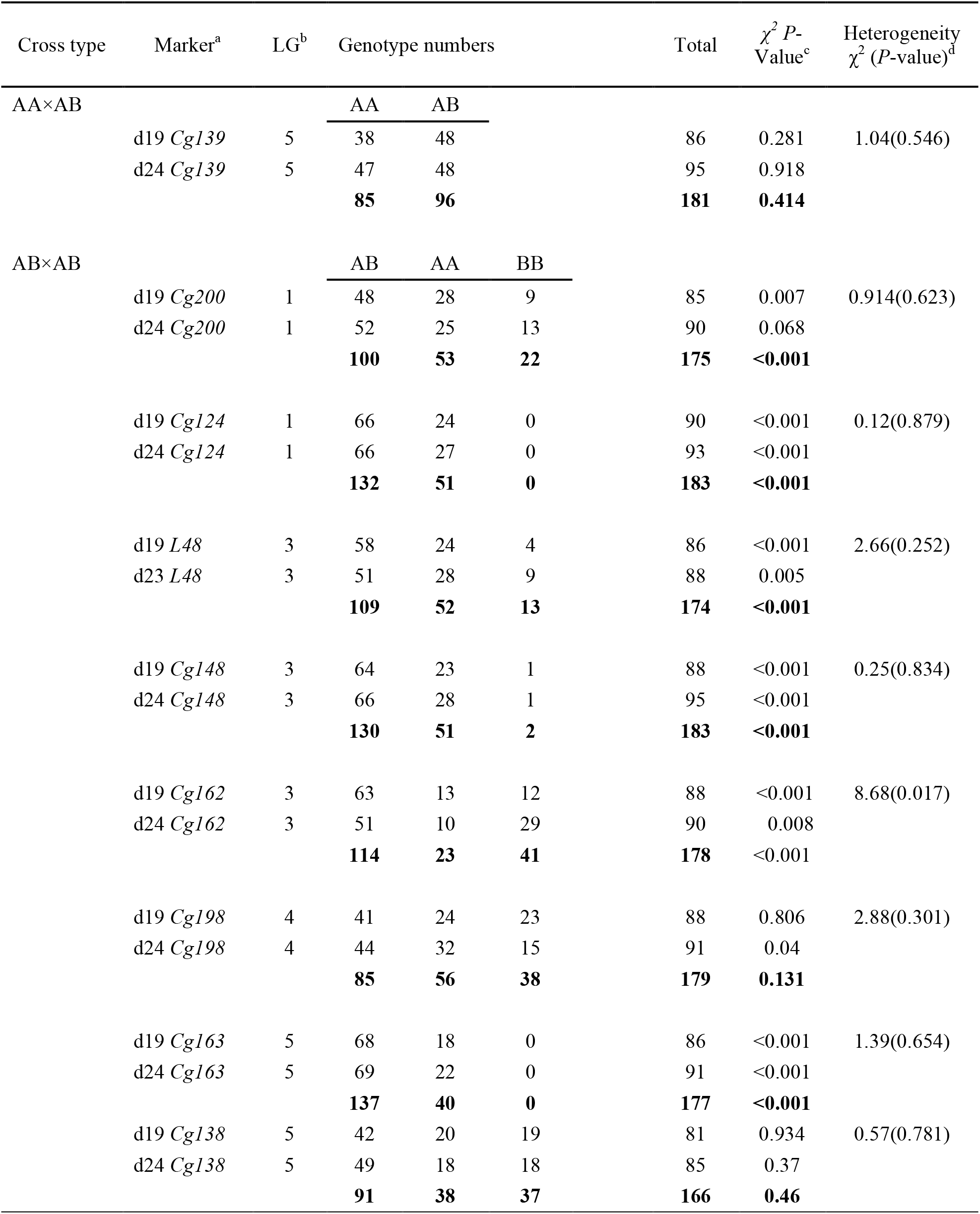
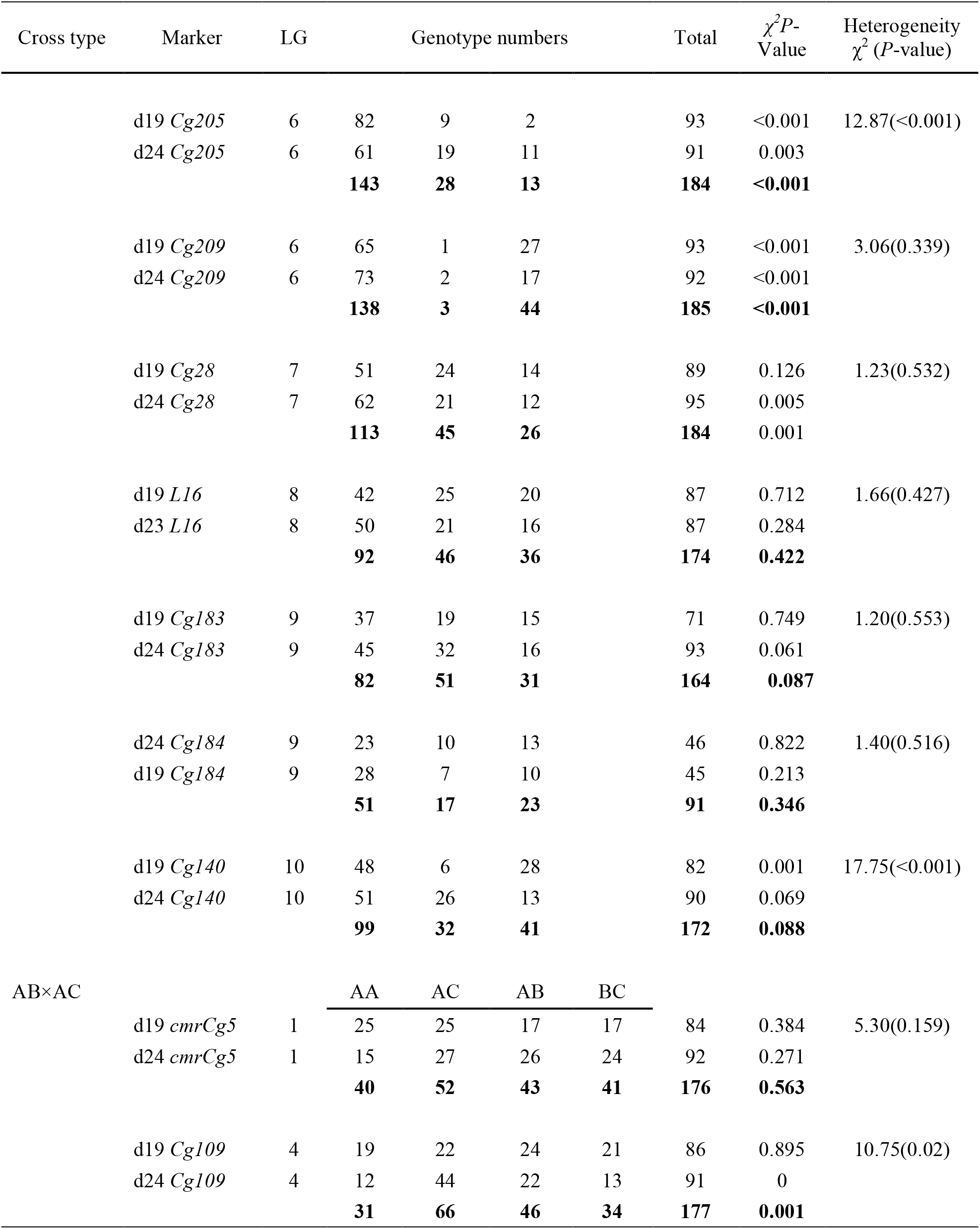
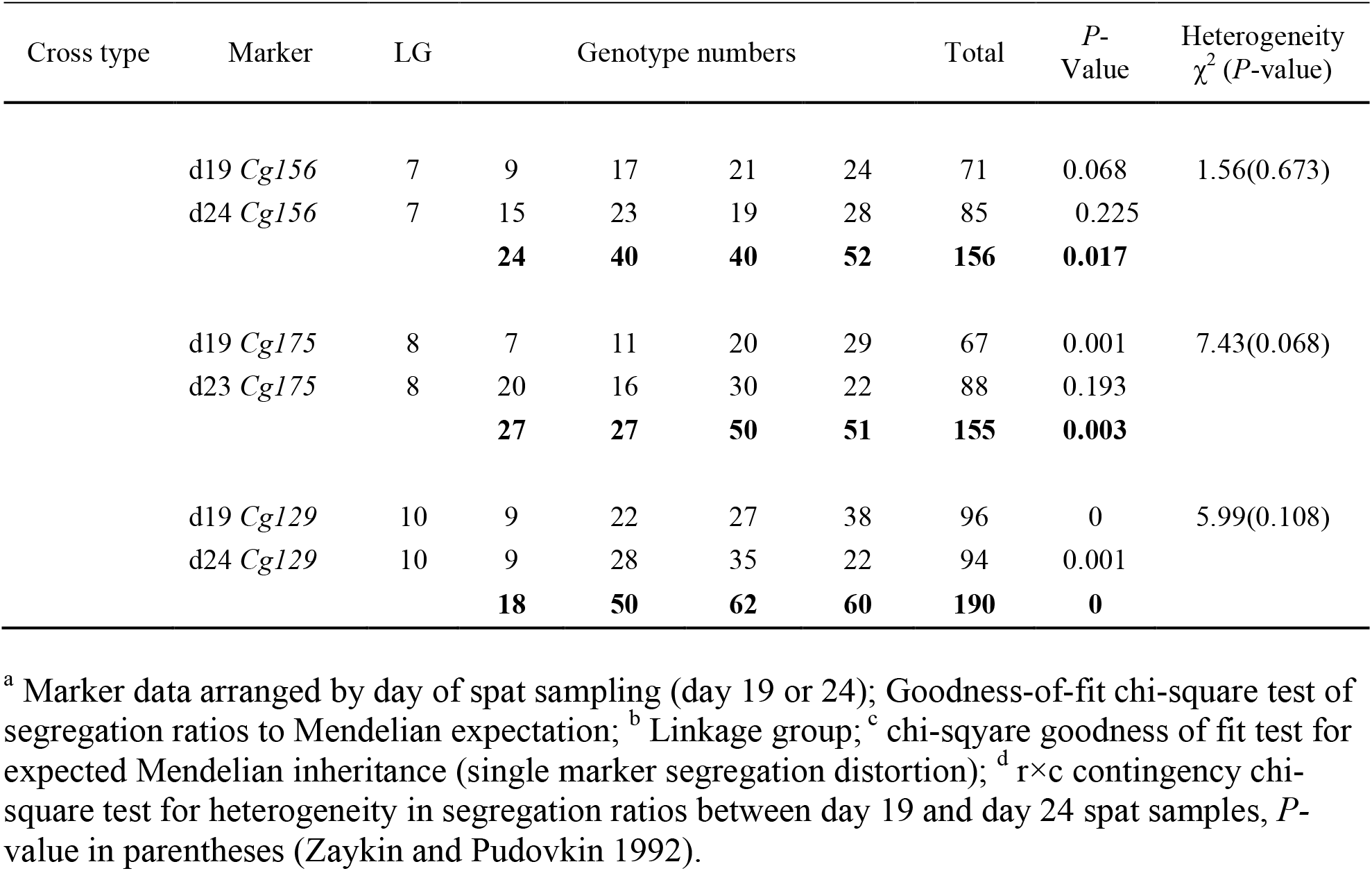
Chi-square tests of genetic heterogeneity between early and late settlement samples

Associations between markers and settlement timing were further assessed in a genome-wide context with quantitative trait locus (QTL) mapping analyses, in which the phenotype, time to settlement, was coded as a binary trait (early or late; settlement on day 19 or 24). The one-dimensional scan identified three QTL on linkage groups 4, 6, and 10 (QTL 1-3; Table 4). The most significant QTL was well above the genome-wide threshold LOD score of 2.99 (QTL 3, LOD=3.794; Table 4) and was located on linkage group 10, closely linked to marker *Cg140*. The other two QTL, one on LG 4 at *Cg109* (QTL 1, LOD=2.567) and one on LG 6 near *Cg205* (QTL 2, LOD=2.714), were slightly below the genome-wide threshold but well above the chromosome-wide thresholds of 2.36 and 1.60 for LG 4 and LG 6, respectively, and are therefore considered suggestive QTL (Table 4). The two-dimensional scan for interacting QTL identified one significant interaction between genomic regions on LG 5 and LG 8 (QTL 4; LG 5 at 10cM, and LG 8 at 40cM, LOD = 5.642), which was above the α = 0.05 genome-wide interaction threshold LOD score of 5.46. Fitting a model for the four identified QTL (from the single QTL scan and the epistasis scan), the ‘drop-one’ analysis of variance (ANOVA) method was used to examine the fit of the model with each QTL included and then dropped in sequential order. In the drop-one analysis, all QTL remained significant except for the interaction QTL, which was marginally significant (*P*=0.057). Results of the ANOVA with four QTL indicated that the model was highly significant (*P*<0.00001), with the four QTL explaining almost 29% of the variance in settlement time; the percent variation explained by the four QTL is an estimate of broad sense heritability for this trait. The strongest effect was at QTL3 (*Cg140*), which alone explained 8.7 % of the variance in settlement timing. Examining the gene effects of the single QTL at the nearest markers, *Cg140* (QTL 3) demonstrated a qualitatively additive pattern; the binomial probability of settlement on day 24 (late settlement) was 0.83, 0.49, and 0.33 for individuals with genotypes *AA, AB,* and *BB*, respectively (Fig. 4). Patterns of gene action at *Cg205* and *Cg109* were less straightforward but appeared to be qualitatively non-additive (Fig. 5a,b).

**Table 4.** QTL and ANOVA model results for settlement timing

| QTL | LG<sup>a</sup> | Position<br>(cM)<sup>b</sup> | Marker<sup>c</sup> | LOD<sup>d</sup> | % Var.<sup>e</sup> | P-Value ($\chi^2$) |
| --- | --- | --- | --- | --- | --- | --- |
| 1 | 4 | 0.0 | *Cg109* | 2.567 | 3.84 | 0.030 |
| 2 | 6 | 1.0 | *Cg205* | 2.714 | 7.085 | 0.001 |
| 3 | 10 | 1.0 | *Cg140* | 3.794 | 8.7 | <0.001 |
|  | 5 | 10.0<sup>f</sup> | -- | -- | 6.66 | 0.210 |
|  | 8 | 40<sup>f</sup> | -- | -- | 5.18 | 0.079 |
| 4 | 5:8 | 10:40 | -- | 5.642 | 6.97 | 0.052 |
| **Full model** |  |  |  |  | **29.9** | **<0.00001** |

<sup>a</sup> linkage group number; <sup>b</sup> position along linkage group, in centimorgans (cM), the colon represents interaction between the two positions for QTL 4; <sup>c</sup> markers closely linked to QTL, if any (within 15 cM); <sup>d</sup> LOD represents the log of the odds (LOD) score for each identified QTL; <sup>e</sup> percent variance explained by the individual QTL or the full model; <sup>f</sup> these genomic positions were not significant QTL in the 1-dimensional scan and, thus, were not assigned numbers; because a significant interaction between them was detected in the two-dimensional scan, R/qtl includes them in the hierarchical ANOVA model by convention, thus their effects are reported.

**Figure 4.**
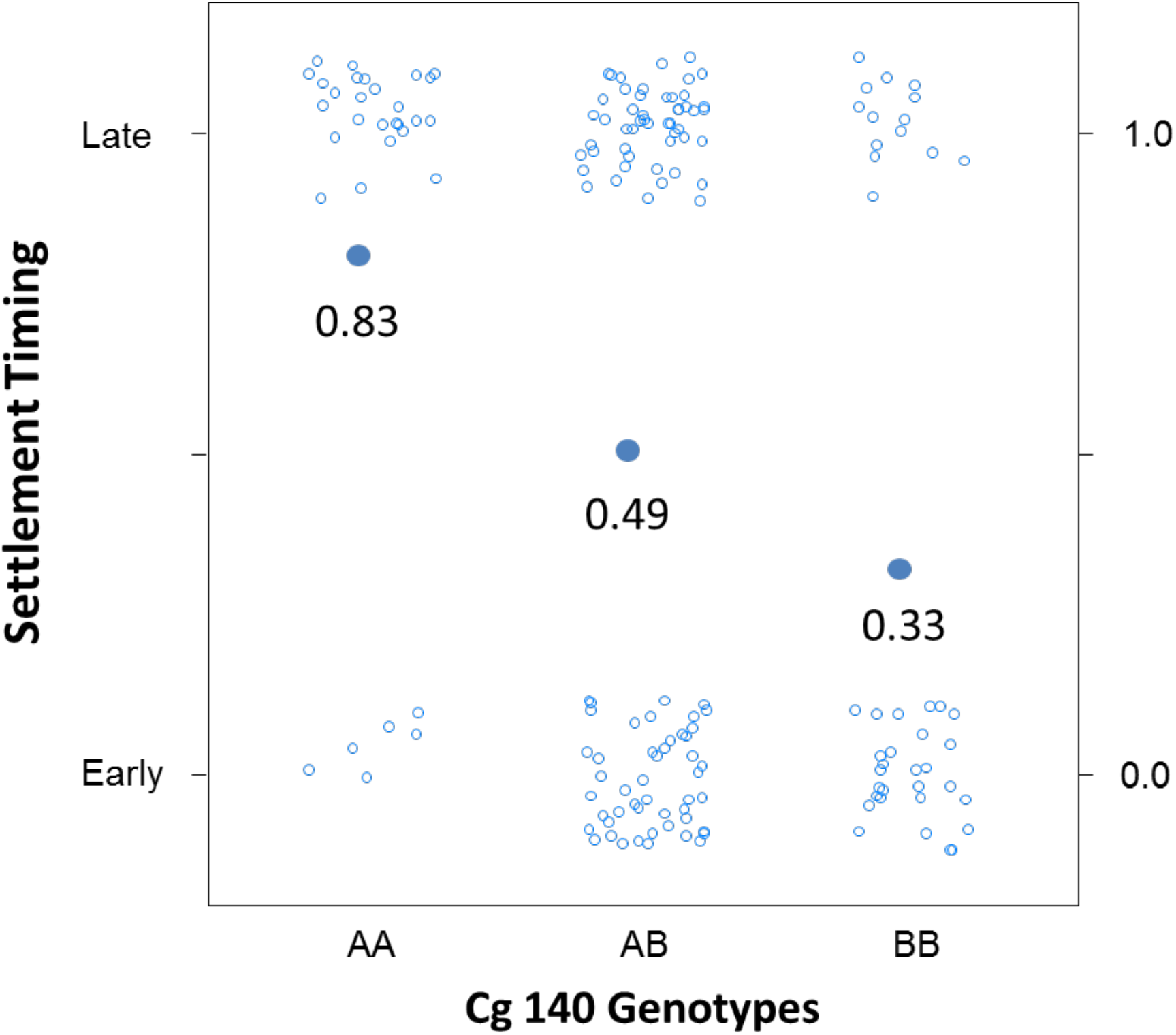
Plot of genotype effects on settlement timing for *Cg140*. Values (0 or 1, corresponding to early and late settlement, respectively) are jittered and represented by open circles. Large blue filled circles represent mean *p* (value of *p* displayed below filled circles), the binomial probability of late settlement for individuals of each genotype, *AA, AB,* and *BB*, which shows an additive pattern of gene effects at this marker.

**Figure 5.**
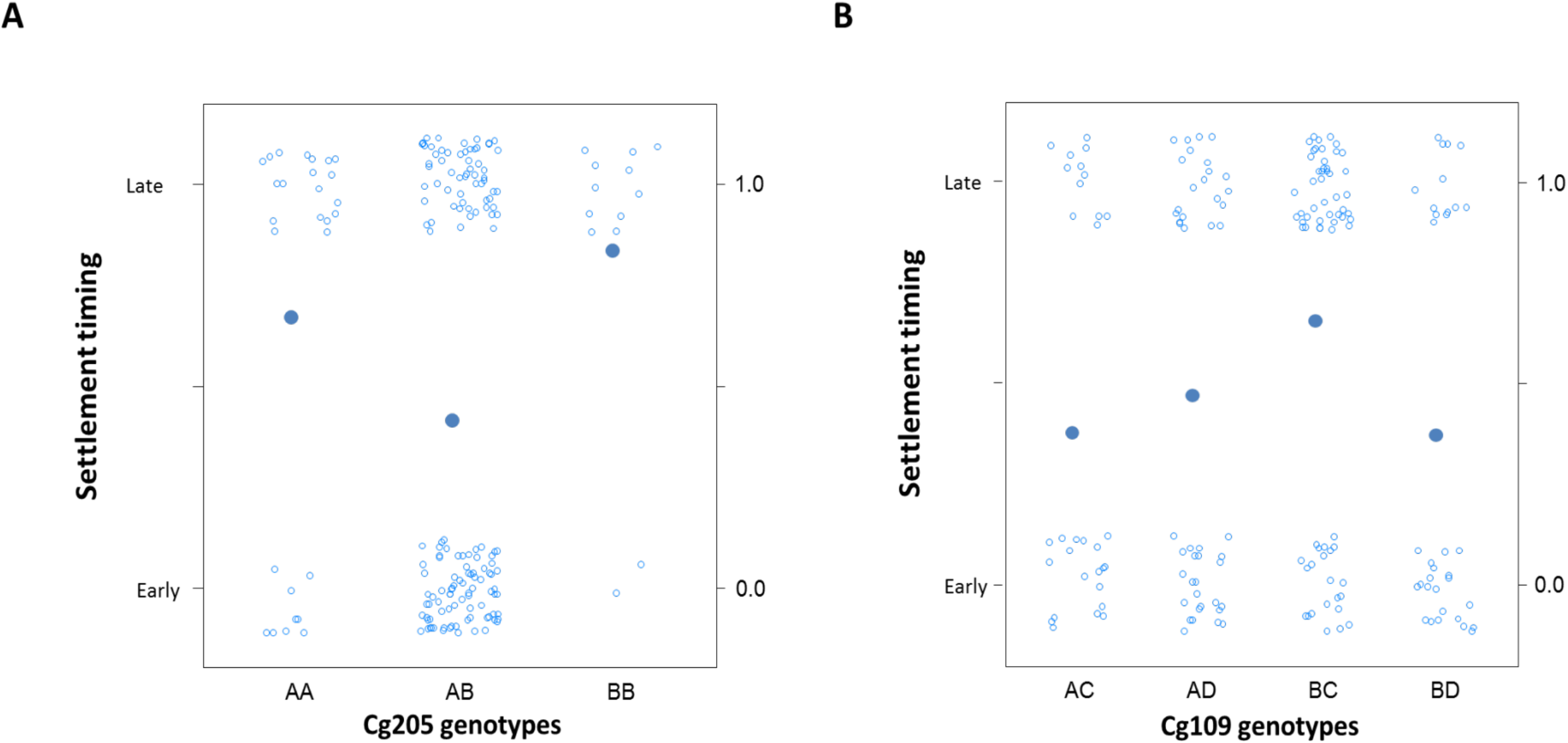
Plot of genotype effects on settlement timing for *Cg205* and *Cg109*. Values (0 or 1, corresponding to early and late settlement, respectively) are jittered and represented by open circles for *Cg205* (panel A) and *Cg209* (panel B). Large blue filled circles represent mean *p*, the binomial probability of late settlement for individuals of each genotype.

## 4. Discussion

In this study, daily sampling of larvae and spat during the settlement of an F_2_ family of the Pacific oyster was carried out to examine, in detail, the patterns of settlement and genotype-dependent mortality during metamorphosis. I first describe the general patterns of settlement and mortality from the observed counts of settlers and larvae, and then turn to the temporal genetic analysis of segregation distortion during metamorphosis, which addresses the biologically relevant hypotheses regarding patterns of mortality during settlement and metamorphosis. Finally, I discuss results of the QTL analysis of settlement timing and the resolution of genetic patterns underlying early vs. late settlement with genotype dependent mortality at metamorphosis.

### 4.1. Settlement patterns and post-settlement mortality

Mortality during settlement was very high, with only 2.8% of late stage larvae surviving as spat at day 60. Compared with survival up to the end of the larval stage (~13%), survival through settlement was almost five-times lower, which confirms settlement and metamorphosis as a critically important life-history transition, during which substantial mortality occurs (*e.g.* Hunt and Sheibling 1996, Plough and Hedgecock 2011, Plough 2012). Survival over the duration of the experiment (early embryo to day 60 spat) was 0.37%, which is consistent with previous reports of substantial early mortality in the Pacific oyster (Bucklin 2003, Plough and Hedgecock 2011, Plough 2012, Plough et al. *in review*). Comparing daily settlement and larval abundance, high rates of mortality in the larval pool appeared to coincide with the peaks of settlement (Fig. 1), suggesting that larval mortality was associated with some aspect of the metamorphic transition: larvae were “trying and dying”. Another striking result was the bimodal distribution in settlement timing: pulses of settlement on days 19 and 24 accounted for 68% of all settlement in this family over the nine-day period. Field-based recruitment studies have shown that settlement during the course of a season is often non-random, with specific peaks of settlement and variable survival over time (*e.g.* Raimondi 1990, Pineda 1994, Balch and Sheibling 2000, Broitman *et al.* 2000, Pineda et al 2006). Temporal variation in settlement in the marine environment could be related to a number of factors including reproductive condition, spawning timing, or oceanographic features such as currents and food availability, none which were variable or relevant in this experiment. Instead, variation in settlement timing, particularly the early and late peaks of settlement, may somehow depend on genetic variability for settlement timing within the family (see below). Only one family was examined in the current study, thus, the observed settlement patterns may not be general; however, a similar bi-modal pattern was observed in a previous experiment (Plough, unpublished data).

Analysis of spat survival 40 days after initial settlement showed conclusively that very little endogenous (genotype-dependent) mortality occurs post-settlement. Essentially, once larvae had completed metamorphosis and appeared as spat on adult shell (time to census was 24 hours or less), individuals survived at a rate of nearly100%. These findings support previous observations of low genotype-dependent morality after metamorphosis (Bucklin 2003, Plough and Hedgecock 2011). Settlement studies of natural populations do, of course, find substantial post-settlement mortality in a variety of marine invertebrates (Hunt and Shiebling 1996, Gosselin and Qian 1997, Pineda et al. 2006) and for oysters in particular (e.g. Kennedy *et al.* 1996, Newell *et al*. 2000). In this experiment, no external sources of mortality, such as predators or environmental stressors were present, thus, these results suggest only that genetic or endogenous causes of mortality do not greatly affect oyster survival once settlement is complete, at least under the laboratory conditions used in this study.

### 4.2. Temporal genetic analysis of mortality during settlement and metamorphosis

At the outset of this study, three hypotheses were proposed to explain the previously observed patterns of substantial genotype-dependent mortality at metamorphosis in inbred crosses of the Pacific oyster: 1) mortality occurs post-settlement, owing to genetic or developmental abnormalities affecting fitness just after the transition to the juvenile stage, 2) mortality occurs during the process of metamorphosis, owing to the expression of deleterious mutations in genes critical for morphogenesis and, 3) mortality occurs prior to metamorphosis, owing to the expression of mutations affecting the development of competency or delaying metamorphosis. The observation of very minimal mortality after settlement and metamorphosis suggests that the expression of genetic load does not greatly affect viability after metamorphosis and, thus, hypothesis one is falsified, at least based on data from the current experiment and our particular culture conditions. However, given the alternative temporal patterns of selection against the two homozygous genotypes at marker *Cg205* during settlement, there is support for the hypotheses that selection is acting at or just preceding metamorphosis (hypotheses two and three). The relatively early and rapid mortality of *AA* individuals during metamorphosis in both the larval and spat samples suggests the expression of a linked, deleterious recessive mutation possibly affecting the morphogenetic transition (hypothesis 2). In contrast, selection against individuals possessing the *BB* genotype appeared to be delayed in the larval samples (high frequency of *BB* larvae on day 26 that never settled in appreciable numbers), which suggests that the *B* allele may be linked to a mutation causing either a delay in metamorphosis or the inability to develop competence and thus begin metamorphosis (hypothesis 3). Of course, it is impossible to explicitly link the observed temporal patterns of selection against each homozygote to specific defects in the morphogenetic or competency pathways, i.e. genes, thus our evaluation of the developmental processes affected is necessarily somewhat qualitative. However, the two different temporal patterns of selection at *Cg205* align well with what we might expect to observe in individuals that fail to initiate metamorphosis (*BB*) or that die during the metamorphic transition (*AA*). Because only a single marker was followed, it is unknown whether mortality related to pre-metamorphic processes (competency or initiation of metamorphosis) or defects in the morphogenetic transition are more typical. Clearly, more markers need to be included across a large number of families to make any conclusive statements about the patterns of metamorphic mortality in Pacific oysters. Overall, with segregation data at *Cg205*, there is support for both hypothesis two and three; endogenous mortality appears to be confined to the larval stages and to metamorphosis, which is again consistent with observations of substantial larval and setting mortality in the hatchery (Plough and Hedgecock 2011, Plough 2012, Plough et al. *in review*). Endogenous mortality caused by the expression of harmful allelic variants may be confined to the early life history stages, because this part of the life cycle is marked by the most substantial developmental changes (multiple larval developmental stages and metamorphosis), while, after the transition to the juvenile stage and associated ecology, few drastic morphological and physiological shifts occur. In other words, most of the “new” genes that are called upon in response to developmental and ecological shifts will already have been “switched on” by the early juvenile phase, and relatively little mortality, owing to large-effect, loss-of-function mutations in critical developmental pathways is expected.

### 4.3. QTL analysis of settlement timing

The finding of significant genetic variance in settlement timing (29% variance explained by 4 QTL) agrees with previous studies by Jackson *et al.* (2005) and Levin et al. (1996), who also found significant genetic (paternal) effects on the development of competency and early settlement in crosses of the vetigastropod *Haliotus spp,* and the polychaete *Streblospio benedici*, respectively. The current study is the first to associate specific genetic markers with settlement timing variation, and even with relatively sparse genomic coverage (21 markers), there is evidence of loci with major effects on variation in this important larval trait. Furthermore, the finding that one of the markers linked to settlement timing QTL showed qualitatively additive patterns of gene action (Cg 140; Fig. 4) suggests that natural or artificial selection could directionally shift mean time to settlement or increase its uniformity (reduced variance), both of which might be valuable breeding improvements for the oyster aquaculture industry (negative correlations with other value traits would need to be examined). Though specific larval behaviors that influence the actual timing or choice of settlement may be associated with the identified QTL, it is possible, or even likely, that other correlated traits could also explain genetic differences between early and late settlement. Time to settlement is essentially a composite larval trait, integrating growth rate at multiple larval stages as well as the attainment of competency, and genetic variance in any of these larval or developmental traits may be reflected in the apparent association between genotype and time to settlement. Indeed, previous studies in the Pacific oyster have shown that hatchery selection for larval growth rate has resulted in a simultaneous decrease in the average time to settlement, suggesting that growth rate and settlement timing may be correlated (Taris *et al.* 2005). Additional evidence from quantitative genetic studies has confirmed that genetic correlations between larval growth rate and time to settlement exist (e.g. Ernand *et al.* 2003). Larval growth rate was not independently measured in this study, so a direct comparison of genetic variance for growth and settlement timing is not possible. Nevertheless, the identification of specific genetic markers that are associated with variance in larval duration is significant, and future QTL-mapping studies employing greater marker densities and measurements of other larval growth-related traits will be better able to characterize loci underlying variation in the larval developmental program.

It is also important to note the possibility that the identification of significant QTL for settlement timing could also be explained by genotype-specific mortality during settlement. For example, the apparent delay of metamorphosis caused by one of the linked mutations at locus *Cg205* (genotype *BB*) could also drive the association of this marker and heterogeneity in settlement timing. Indeed, two of the three identified QTL for settlement timing were also associated with deleterious mutations acting during metamorphosis *(Cg205* and *Cg109*), which supports this hypothesis. In these cases, it is difficult to distinguish between the action of genotype-dependent mortality and alleles that directly alter settlement behavior and ultimately the timing of settlement. On the other hand, the most significant QTL identified in this study appears to be independent of genotype-dependent mortality at metamorphosis. *Cg140* (closely linked to QTL 3) did not experience significant genotype dependent mortality (chi-square goodness-of-fit test in the final, pooled, day 60 spat sample, *P*=0.088; Table 1), which suggests it is not closely linked to a viability mutation. Thirty centiMorgans (cM) away on the same linkage group (LG 10), *Cg129* is significantly distorted, but prior to metamorphosis (*P*<0.0001, goodness-of-fit chi-square test for the day 18 larval sample; Table 1), demonstrating that while there is genotype-dependent mortality on this linkage group, it is pre-metamorphic and does not appear to influence genotype frequencies at *Cg140*. Thus, genotype-dependent mortality during metamorphosis cannot explain genotype frequency differences between early and late settlement for the QTL linked to *Cg140*. The epistatic QTL identified in this study (QTL 4) is also not likely to be explained by genotype-dependent mortality at metamorphosis, as interactions among loci linked to deleterious mutations (viability loci) were virtually absent in all previous crosses examined (Bucklin 2003, Plough and Hedgecock 2011, Plough 2012). Overall, the confounding effects of genotype-dependent mortality (at metamorphosis) on the identification of QTL for settlement timing warrant serious consideration but likely do not completely account for the QTL results in this particular experiment.

## Acknowledgements

The author wishes to thank D. Hedgecock and S. Edmands for helpful comments and suggestions on previous versions of the manuscript. This work was funded in part by an Oakley Fellowship and a Rose Hills foundation fellowship to L. Plough.

